# Exposure to the gut microbiota from cigarette smoke-exposed mice exacerbates cigarette smoke extract-induced inflammation in zebrafish larvae

**DOI:** 10.1101/2021.09.20.461170

**Authors:** Simone Morris, Kathryn Wright, Vamshikrishna Malyla, Warwick J Britton, Philip M Hansbro, Pradeep Manuneedhi Cholan, Stefan H Oehlers

## Abstract

Cigarette smoke (CS)-induced inflammation leads to a range of diseases including chronic obstructive pulmonary disease and cancer. The gut microbiota is a major modifying environmental factor that determine the severity of cigarette smoke-induced pathology. Microbiomes and metabolites from CS-exposed mice exacerbate lung inflammation via the gut-lung axis of shared mucosal immunity in mice but these systems are expensive to establish and analyse. Zebrafish embryos and larvae have been used to model the effects of cigarette smoking on a range of physiological processes and offer an amenable platform for screening modifiers of cigarette smoke-induced pathologies with key features of low cost and rapid visual readouts. Here we exposed zebrafish larvae to cigarette smoke extract (CSE) and characterised a CSE-induced leukocytic inflammatory phenotype with increased neutrophilic and macrophage inflammation in the gut. The CSE-induced phenotype was exacerbated by co-exposure to microbiota from the faeces of CS-exposed mice, but not control mice. Microbiota could be recovered from the gut of zebrafish and studied in isolation in a screening setting. This demonstrates the utility of the zebrafish-CSE exposure platform for identifying environmental modifiers of cigarette smoking-associated pathology and demonstrates that the CS-exposed mouse gut microbiota potentiates the inflammatory effects of CSE across host species.

## Introduction

Smoking tobacco is one of the greatest public health challenges, accounting for more than 8 million deaths worldwide, per year (1). It ranks as the third leading risk factor for illness and death, including from diseases such as cancers and inflammatory disorders like inflammatory bowel disease (IBD) (1–4). Cigarette smoke (CS) contains a multitude of toxic compounds affecting cell function, foetal development, gut function and microbiome, neural activity, inflammation, and lung cancer (3, 5–12).

The toxicity of CS has further been demonstrated to alter the microbiome of smokers, resulting in respiratory and gut microbiome dysbiosis, which in turn is associated with increased peptic ulcers, IBD and gastrointestinal cancers via the gut-lung axis of shared mucosal immunity (3, 4, 6, 11, 13). This is suggested to occur through alterations to the immune system (5), mucin production (6) or through direct anti-microbial effects of the toxins within the smoke (14). Shifts in gut microbiome and the profile of gut metabolites and pathology are associated with chronic obstructive pulmonary disease (COPD) in humans and CS-induced emphysema in mice (4, 10, 11, 15, 16). Microbiomes and metabolites associated with COPD and CS exacerbate lung inflammation via the gut-lung axis of shared mucosal immunity contributing to lung pathology in mammals (4, 11, 17).

Zebrafish have been used to study the developmental defects caused by CS exposure. These studies demonstrated stunted embryonic development upon exposure to cigarette smoke extract (CSE) and smoke condensate (18–20). Exposure of adult zebrafish to CSE reduced the capacity for fin regeneration (19), as well as behavioural abnormalities in response to nicotine (7). A study by Progatzky et al. further modelled the respiratory effects of CS using adult zebrafish gills demonstrating increased pro-inflammatory cytokine production and reduced immune cell recruitment (21). Massarsky et al. used total particulate matter (TPM) from cigarettes to model the effects of CS in zebrafish, and showed cardiac malfunction and defects in brain vascularisation (22) as well as behavioural alterations (23).

Thus far, the effect of CS exposure on immune function has been unexplored in zebrafish embryos and larvae (19, 21). This transparent stage of development provides the opportunity to investigate the inflammatory effects of CSE by live imaging of fluorescently tagged leukocytes, and is amenable to simplified experimental manipulation by gene editing or treatment with small molecules by immersion (24). Here we define the pro-inflammatory effects of CSE on larval zebrafish and find a transferrable effect of mammalian CS-associated microbiota on CSE-induced inflammation in larval zebrafish.

## Methods

### Zebrafish husbandry

Zebrafish embryos were obtained through natural spawning (Sydney Local Health District Animal Welfare Committee Approval 17-036). The transgenic strains used to visualise leukocytes were *Tg(lyzC:DsRed)*^*nz50*^, where neutrophils are labelled by DsRed2 expression (25), and *Tg(mfap4:turquoise)*^*xt27*^, where macrophages are labelled by Turquoise2 (26). Embryos were raised in E3 media supplemented with methylene blue in a dark incubator at 28°C until 1 day post fertilisation (dpf). Cleaned 1 dpf embryos were transferred to E3 without methylene blue and reared (dark incubator, 28°C) until used.

Zebrafish larvae were anaesthetised in 250 μg/mL MS-222 (Tricaine, Sigma) and euthanised either by fixation in 4% paraformaldehyde, lysis in Trizol, or disposed of in bleach as per biosafety protocols.

### Preparation of CSE

Cigarette smoke extract (CSE) was prepared by using a custom-built smoking device where reference cigarette 3R4F (Kentucky Tobacco Research & Development Centre) was lit using a lighter and CS was bubbled directly in to 10 mL of PBS until the whole cigarette was exhausted. This 10 mL of CSE was filtered using a 0.22-micron filter to remove CS-derived debris. This preparation of the smoke of one cigarette per 10 mL PBS is considered 100% CSE and was diluted as indicated. CSE was prepared fresh before treatment.

### Exposure of zebrafish to CSE

Embryos/larvae were exposed to CSE at a density of approximately 1 larva per mL of E3 in plastic Petri dishes at the ages indicated, either 3 or 5 dpf, and maintained in a dark incubator at 28°C. The stock solution of 100% CSE in PBS was diluted directly into E3 zebrafish media to a final concentration of 1%, 2%, or 3%. Filter sterilised PBS was used as vehicle control.

### Zebrafish behavioural analysis

Zebrafish larvae were transferred to a 24-well plate with 1 embryo per well in their CSE-supplemented E3 media. The plate was transferred to a ZebraBox (Viewpoint) and locomotion was recorded for 15 minutes. Video files were analysed using EthoVision XT 11 (Noldus) using the following parameters: sensitivity 110, video pixel smoothing low, subject size 4-4065. Locomotion analysis was calculated as the total distance travelled (mm) of individual larva over the recording window.

### Tail transection assays

Caudal fin amputations were performed on 5 dpf larvae. Larvae were anesthetised with 2.5% (v/v) ethyl-3-aminobenzoate methanesulfonate (tricaine) (Sigma, E10521), wounded posterior to the notochord using a sterile scalpel and kept in a 28°C incubator to recover as previously described (27). Wounded larvae were imaged at 6 hours post wounding.

### Imaging and image analysis

Larvae were imaged using a Leica M205FA fluorescent microscope. ImageJ software was used to quantify fluorescent pixel counts as previously described (27).

### Generation of microbiome-depleted zebrafish embryos

Microbiome-depleted zebrafish were created and maintained as previously described (28). Briefly, freshly laid embryos were rinsed with 0.003% v/v bleach in sterile E3 and rinsed 3 times with sterile E3. Bleached embryos were raised in sterile E3 supplemented with 50 μg/mL ampicillin (Sigma), 5 μg/mL kanamycin (Sigma) and 250 ng/mL amphotericin B (Sigma) in sterile tissue culture flasks. Dead embryos and chorions were aseptically removed at 1 and 3 dpf, respectively.

### Generation of mouse faecal microbiota specimens

Mice were housed at the Centenary Institute (Sydney Local Health District Animal Welfare Committee Approval 2020-003). C57BL/6J mice were housed in a pathogen-free and temperature-controlled environment, with 12 hours of light and 12 hours of darkness, and free access to food and water. Mice were smoked with 12x 3R4F cigarettes for 75 minutes, twice a day, 5 days per week for 10 weeks by using a custom-designed and purpose-built nose-only, directed flow inhalation and smoke-exposure system (CH Technologies, NJ) (10, 29). This is representative of a pack-a-day smoker (2).

Faecal pellets were collected from mice that were housed in different cages. Individual faecal pellets were collected into sterile 1.7 mL microcentrifuge tubes and homogenised in 1 mL of sterile E3 by pipetting. Homogenised specimens were centrifuged (500 *xg*, 2 minutes) to sediment fibrous material and supernatant were collected then pooled. Pooled supernatants were supplemented with glycerol to a final concentration of 25% v/v, aliquoted, and frozen at −80°C before use.

Mice were not anaesthetised for these procedures and were used for other experiments as per Approval 2020-003 where they were eventually sacrificed by CO2 asphyxiation.

### Exposure of zebrafish larvae to mouse faecal microbiota

Microbiome-depleted zebrafish were rinsed at 3 dpf with sterile E3, without antibiotic and amphotericin B supplementation, and colonised by the addition of 200 μL thawed faecal homogenate supernatant into 30 mL sterile E3 delivering a final concentration of ~5×10^4^ CFU per mL. Enumeration of culturable bacterial load was carried out by culture under aerobic conditions at 28°C on LB agar (Thermofisher).

### Detection of gene expression

Groups of at least 10 larvae were homogenised in Trizol (Thermofisher) and processed for RNA extraction as per manufacturer’s instructions. cDNA was reverse transcribed using the Applied Biosystems High Capacity cDNA synthesis kit (Thermofisher) from 2 μg of total RNA. Quantitative PCR was carried out on a CFX Connect machine (Biorad) with Powerup SYBR reagent (Thermofisher), primer sets were (5’-3’): il1b Fw ATCAAACCCCAATCCACAGAGT, Rv GGCACTGAAGACACCACGTT; 18s Fw TCGCTAGTTGGCATCGTTTATG, Rv CGGAGGTTCGAAGACGATCA.

### Statistical analysis

Statistical testing was carried out by ANOVA with post hoc multiple comparisons as appropriate using GraphPad Prism 9 (GraphPad). Data are expressed as mean ± SD, calculated P values are provided in graphs. Outliers were excluded using a 1% robust regression and outlier removal (ROUT) test. Every datapoint represents a single embryo unless otherwise noted as for qPCR analyses.

## Results

### Chronic exposure to CSE is toxic to zebrafish post-hatching larvae

To establish a survivable dose of CSE that did not cause gross pathology, we exposed 3 dpf zebrafish larvae to a range of CSE doses and monitored survival. We found exposure at up to 2% CSE was tolerated by zebrafish larvae (Figure 1A), with no overt changes to morphology (Figure 1B). Exposure to 3% CSE caused significant mortality within 1 day post exposure.

**Figure 1:**
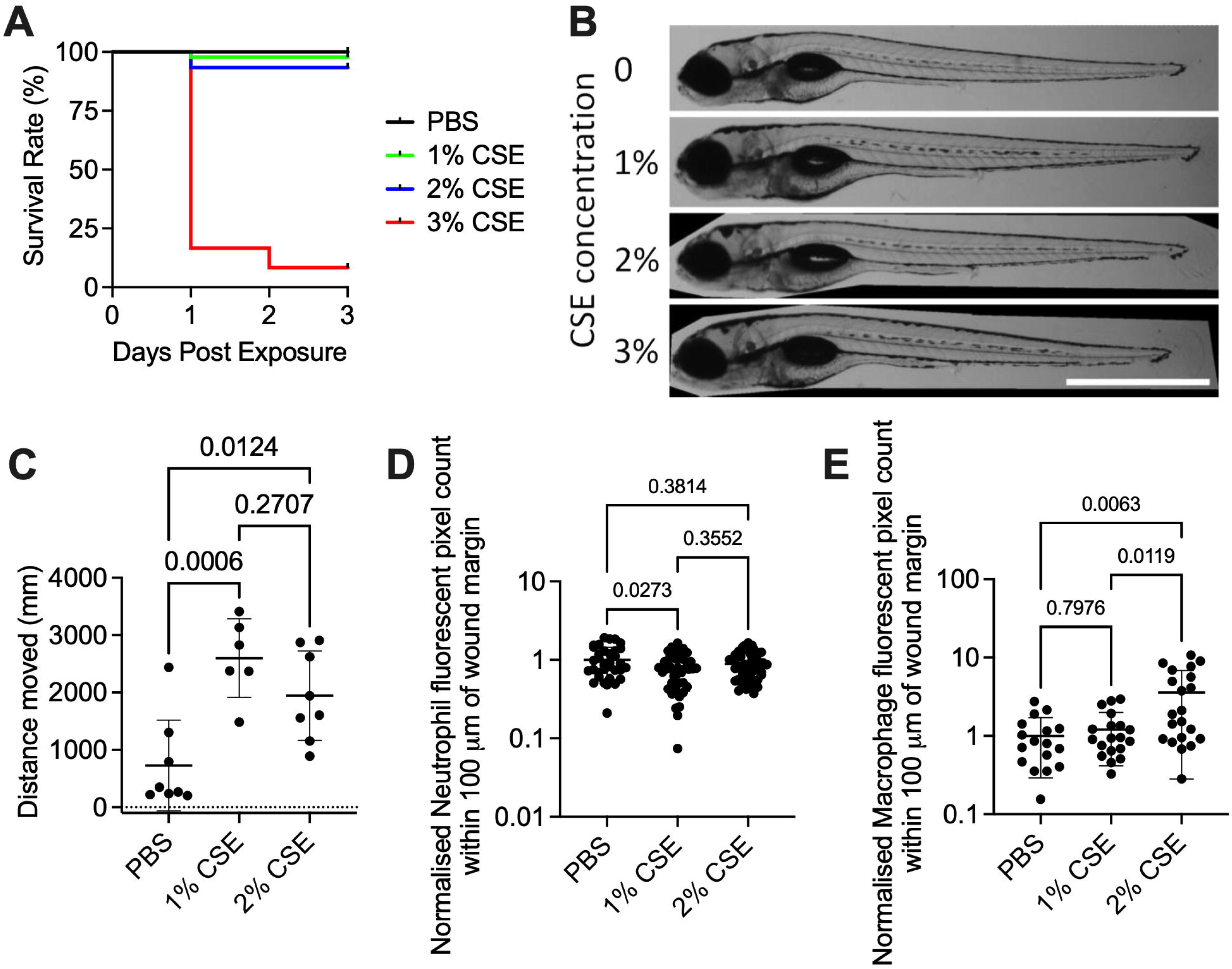
Cigarette smoke extract is toxic to zebrafish larvae, and causes increased embryo motility, and greater macrophage influx into wounds. A. Survival of 3 dpf larvae after exposure to CSE. Individual Log-rank comparison scores: P(0% vs 1%) = 0.3173, P(0% vs 2%) = 0.0798, P(0% vs 3%) <0.0001, P(1 vs 3) <0.0001, P(2% vs 3%) <0.0001. Starting n per group: 0 = 45 larvae, 1% = 45 larvae, 2% = 45 larvae, 3% = 36 larvae. B. Representative images of 5 dpf larvae exposed to CSE from 3 dpf. Scale bar = 1 mm. C. Total distance moved in 15 minutes by 5 dpf larvae exposed to CSE from 3 dpf. Statistical comparison by ANOVA, data are representative of two biological replicates. D. Quantification of neutrophil fluorescent pixel area within 100 μm of the wound margin at 6 hpw in 5 dpf larvae exposed to CSE from 3 dpf. Statistical comparison by ANOVA, data are normalised and combined from two biological replicates. E. Quantification of macrophage fluorescent pixel area within 100 μm of the wound margin at 6 hpw in 5 dpf larvae exposed to CSE at doses indicated from 3 dpf. Statistical comparison by ANOVA, data are normalised and combined from two biological replicates.

To verify that the CSE was bioactive at 1% and 2% concentrations, we tracked larval motility to detect nicotine-induced increases in motility at 5 dpf, or 2 days post exposure (30). Exposure to 1% or 2% CSE increased the total distance travelled of zebrafish larvae compared to mock exposed larvae in a 15 minute recording period indicating components of the CSE were still bioactive after 2 days of immersion exposure in E3 media (Figure 1C).

We next used *Tg(lyzC:DsRed)*^*nz50*^ larvae to visualise neutrophils and *Tg(mfap4:turquoise)*^*xt27*^ to visualise macrophages (25, 26). Immune cell fluorescent area was used as a proxy to estimate changes to immune cell number by non-invasive live imaging. We performed tail transection on 5 dpf larvae that had been exposed to CSE from 3 dpf and assessed leukocyte recruitment to the wound at 6 hours post wounding. While neutrophil recruitment was slightly but statistically significantly reduced by 1% CSE, there was no effect of 2% CSE exposure compared to vehicle control and overlap between neutrophil recruitment in the 1% and 2% CSE exposure groups (Figure 1D). There was significantly more macrophage recruitment to wounds in 2% CSE-exposed larvae compared to control and 1% CSE larvae (Figure 1E).

### Acute exposure to CSE induces myelopoiesis in zebrafish larvae

To examine the acute effects of CSE exposure on larvae after major organogenesis is complete, we exposed 5 dpf larvae to CSE for 24 hours. Immersion in 1% and 2% CSE caused a gradated increase in total neutrophil fluorescent area in zebrafish larvae compared to PBS vehicle (Figure 2A and 2B). This inflammatory effect of CSE appeared stronger on total macrophage fluorescent area with a statistically significant increase in total macrophages in 2% CSE-exposed compared to control larvae (Figure 2C and 2D).

**Figure 2:**
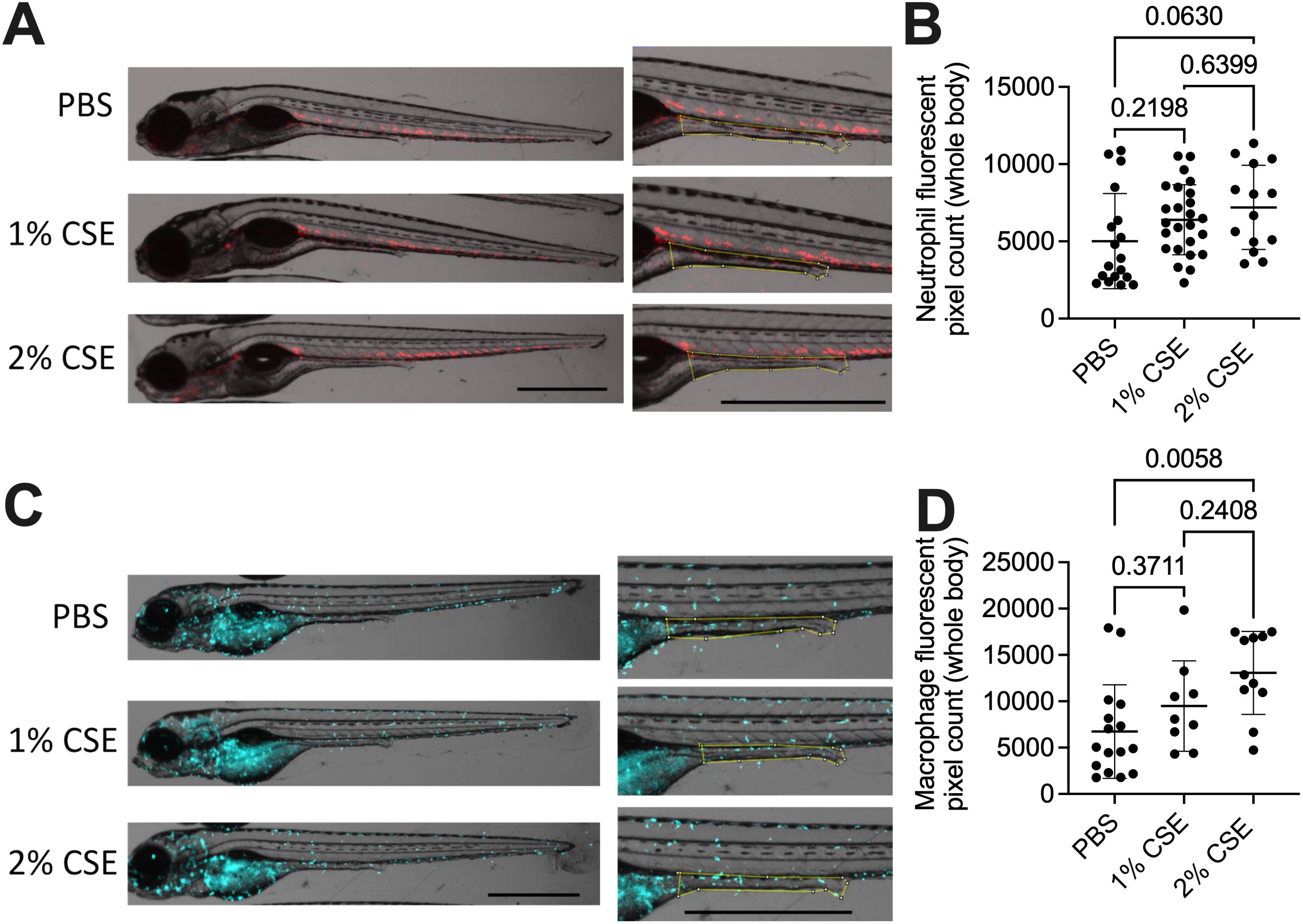
Acute exposure to CSE induces myelopoiesis in zebrafish larvae. A. Representative images of 6 dpf *Tg(lyzC:DsRed)*^*nz50*^ larvae exposed to CSE for 24 hours from 5 dpf. B. Quantification of total neutrophil fluorescent pixel area in 6 dpf larva exposed to CSE for 24 hours from 5 dpf. C. Representative images of 6 dpf *Tg(mfap4:turquoise)*^*xt27*^ larvae exposed to CSE for 24 hours from 5 dpf. D. Quantification of total macrophage fluorescent pixel area in 6 dpf larva exposed to CSE for 24 hours from 5 dpf. A & C: Scale bars = 1 mm. Bounded area in insets are used to measure gut leukocytes in Figure 3 onwards. B & D: Statistical comparison by ANOVA, data are representative of two biological replicates.

### Acute exposure to CSE induces an inflammatory response in the gut of zebrafish larvae

The effects that we observed suggested that CSE induced an inflammatory haematopoietic response. Since the larval midgut and hindgut is an important site of immune response to chemical irritants, we next quantified leukocyte recruitment to the larval intestine of 6 dpf larvae that had been exposed to 2% CSE from 5 dpf (31). Leukocyte fluorescence was increased in the gut compared to either control or 1% CSE-exposed larvae (Figure 3A and 3B). However, the estimation of gut macrophages by fluorescent pixel count in our *Tg(mfap4:turquoise)*^*xt27*^ larvae was potentially confounded by luminal autofluorescence that was more noticeable in CSE-exposed larvae (Figure 2B).

**Figure 3.**
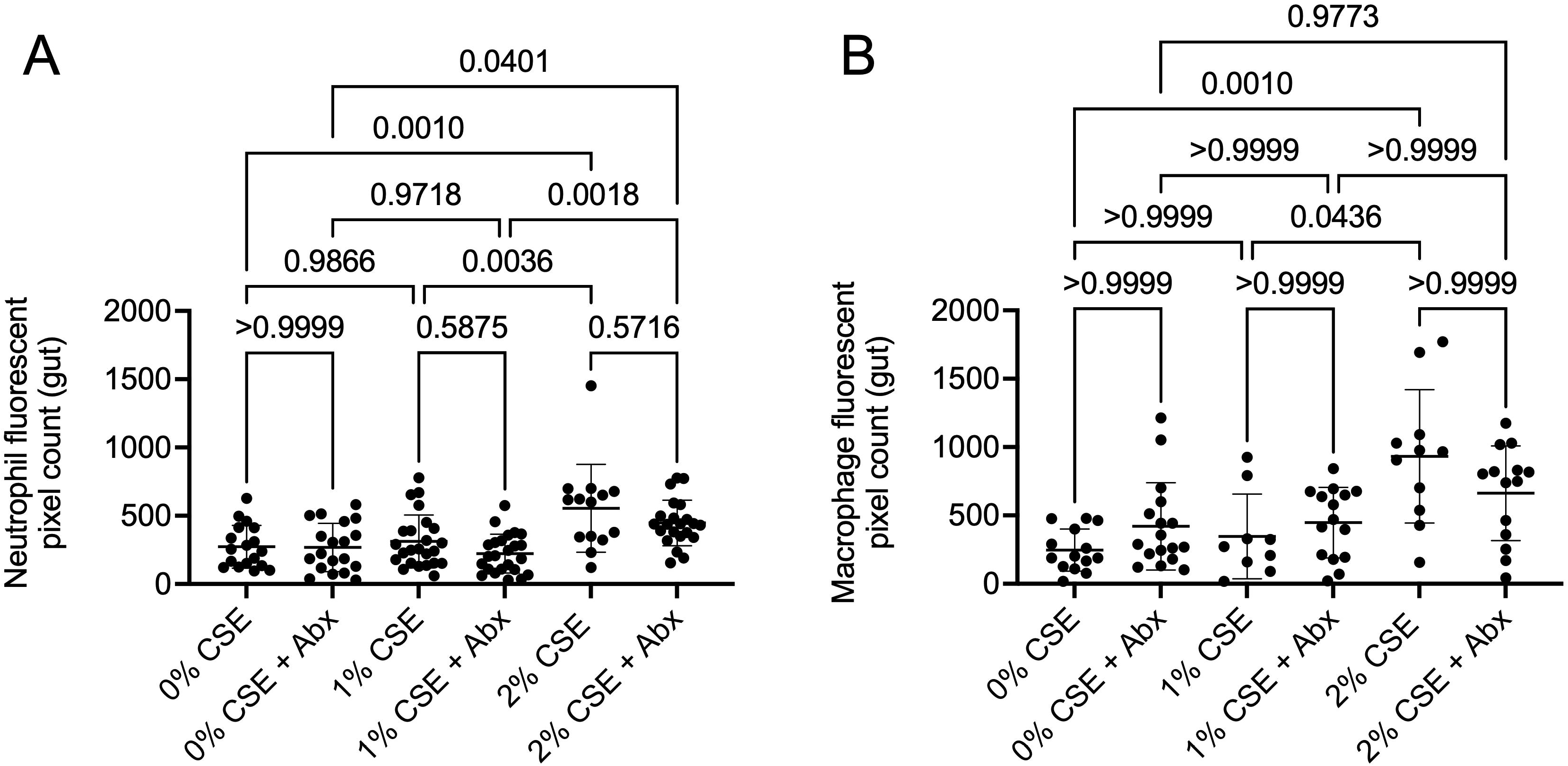
Acute exposure to CSE induces an inflammatory response in the gut of zebrafish larvae. A. Quantification of gut neutrophil fluorescent pixel area in 6 dpf larvae treated with broad spectrum antibiotics (Abx) ampicillin and kanamycin and exposed to CSE from 5 dpf. B. Quantification of gut macrophage fluorescent pixel area in 6 dpf larvae treated with broad spectrum antibiotics (Abx) ampicillin and kanamycin and exposed to CSE from 5 dpf. Statistical comparisons by ANOVA, data are representative of two biological replicates.

The enhanced effects of CSE on innate immune cell numbers in the larval gut led us to hypothesise that the microbiota may interact with CSE to drive inflammation at zebrafish larval mucosal surfaces as observed with other chemical irritants (31, 32). Thus, we next examined the effect of CSE and the role of changes in the endogenous microbiota by depleting bacteria with broad spectrum antibiotics prior to CSE exposure from 5 to 6 dpf. Antibiotic treatment did not affect the ability of 2% CSE to increase neutrophil or macrophage fluorescence in the larval gut suggesting that the natural gut microbiota is not necessary to drive the acute inflammatory effects of 2% CSE (Figure 3A and 3B).

### Microbiota from CS-exposed mice increases the neutrophilic inflammatory effects of CSE

We next assessed if the microbiota of chronically CS-exposed mice could impart increases in sensitivity to CSE exposure by exposing 3 dpf microbiota-depleted larval zebrafish to faecal homogenate from mice exposed to CS/air controls for 10 weeks and CSE. We used the suboptimal concentration of 1% CSE, which did not induce intestinal inflammation in our acute exposure assay, to detect enhancement of inflammation by microbiota derived from the faeces of CS-exposed mice.

The total count of neutrophil fluorescence was unaffected in 5 dpf larvae immersed with microbiota from CS- or normal air-exposed control mice, and the addition of 1% CSE (Figure 4A and 4B).

**Figure 4.**
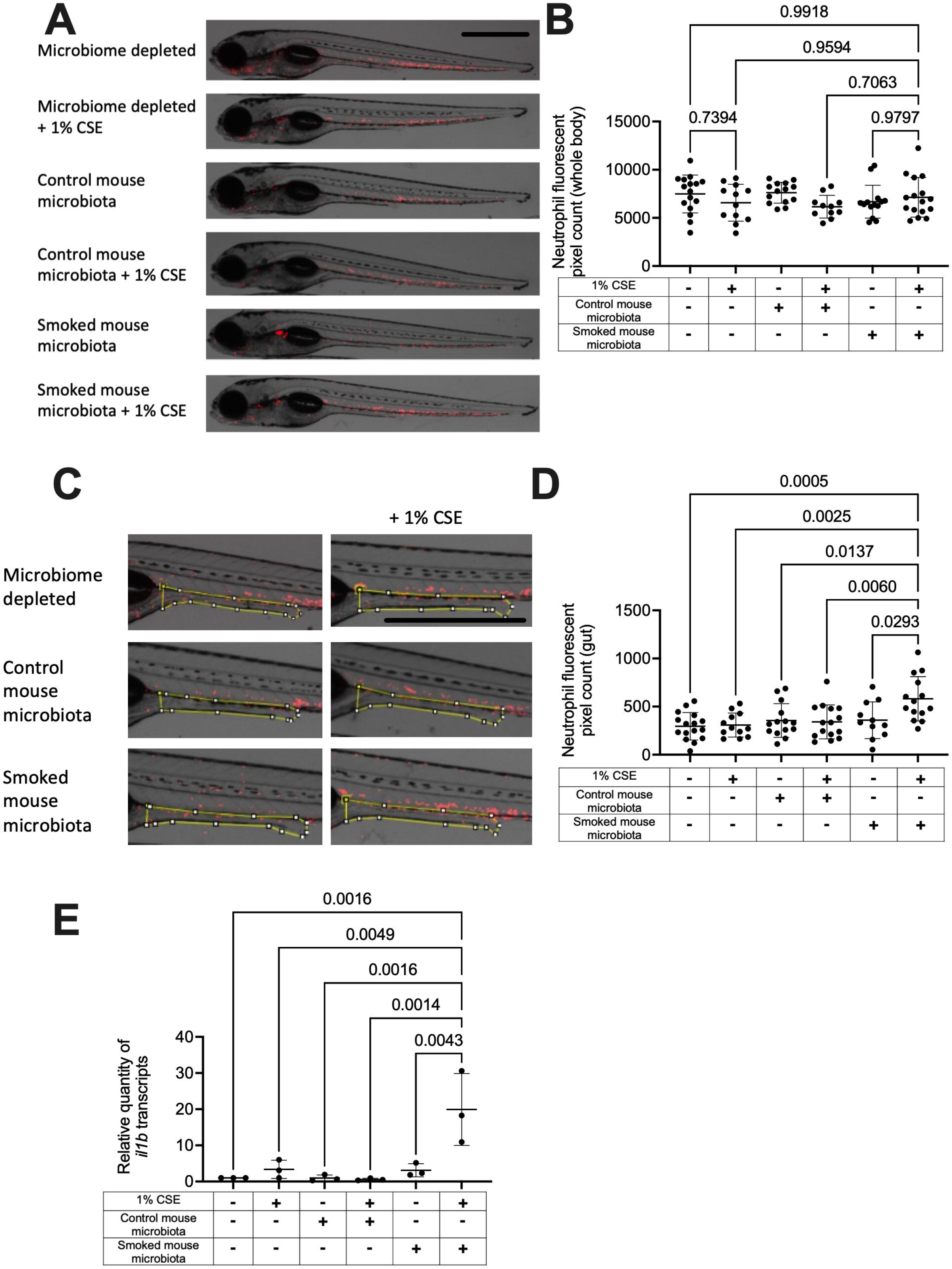
CS-exposed mouse microbiota increases the neutrophilic inflammatory effects of CSE. A. Representative images of 5 dpf microbiota-depleted *Tg(lyzC:DsRed)*^*nz50*^ larvae exposed to combinations of CS- or control normal air-exposed mouse microbiota with or without 1% CSE from 3 dpf. B. Quantification of total neutrophil fluorescent pixel area per larva exposed to combinations of CS- or control normal air-exposed mouse microbiota with or without 1% CSE from 3 dpf. Statistical comparison by ANOVA, data are representative of two biological replicates. C. Representative images of the posterior guts of microbiome depleted *Tg(lyzC:DsRed)*^*nz50*^ larvae exposed to combinations of CS- or control normal air-exposed mouse microbiota with or without 1% CSE from 3 dpf. Area used for analysis demarcated by yellow box. D. Quantification of gut neutrophil fluorescent pixel area per larva exposed to combinations of CS- or control normal air-exposed mouse microbiota with or without 1% CSE from 3 dpf. Statistical comparison by ANOVA, data are representative of two biological replicates. E. Quantification of *il1b* transcripts by qPCR in larvae exposed to combinations of CS- or control normal air-axposed mouse microbiota with or without 1% CSE from 3 dpf. Statistical comparison by ANOVA, each data point represents a biological replicate with at least 10 larvae per group. Scale bars = 1 mm.

Exposure to microbiota from CS-exposed or control mice did not affect neutrophil fluorescence in the gut of wild-type 5 dpf larvae compared to germ-free reared larvae (Figure 4C and 4D). Addition of 1% CSE did not affect neutrophil fluorescence in the gut in 5 dpf larvae raised microbiota-depleted or in those exposed to microbiota from control non-smoked mice. However, 5 dpf larvae that had been exposed to microbiota from smoked mice and 1% CSE had significantly more neutrophil fluorescence in the gut than any of the control groups. This pattern was replicated by *il1b* transcription levels quantified by RT-qPCR (Figure 4E).

### CS-exposed mouse microbiota increases the macrophage inflammatory effects of CSE

Analysis of *Tg(mfap4:turquoise)*^*xt27*^ macrophage reporter larvae revealed increased total macrophage fluorescence in 5 dpf larvae that that had been exposed to microbiota from CS-exposed mice and 1% CSE compared to any of the control groups, except for the air-exposed mouse microbiota and 1% CSE control group (Figure 5A and 5B). Quantification of gut macrophage fluorescence was again confounded by luminal autofluorescence in CSE-exposed larvae, appearing to be elevated in all CSE-exposed groups (Figure 5C and 5D).

**Figure 5.**
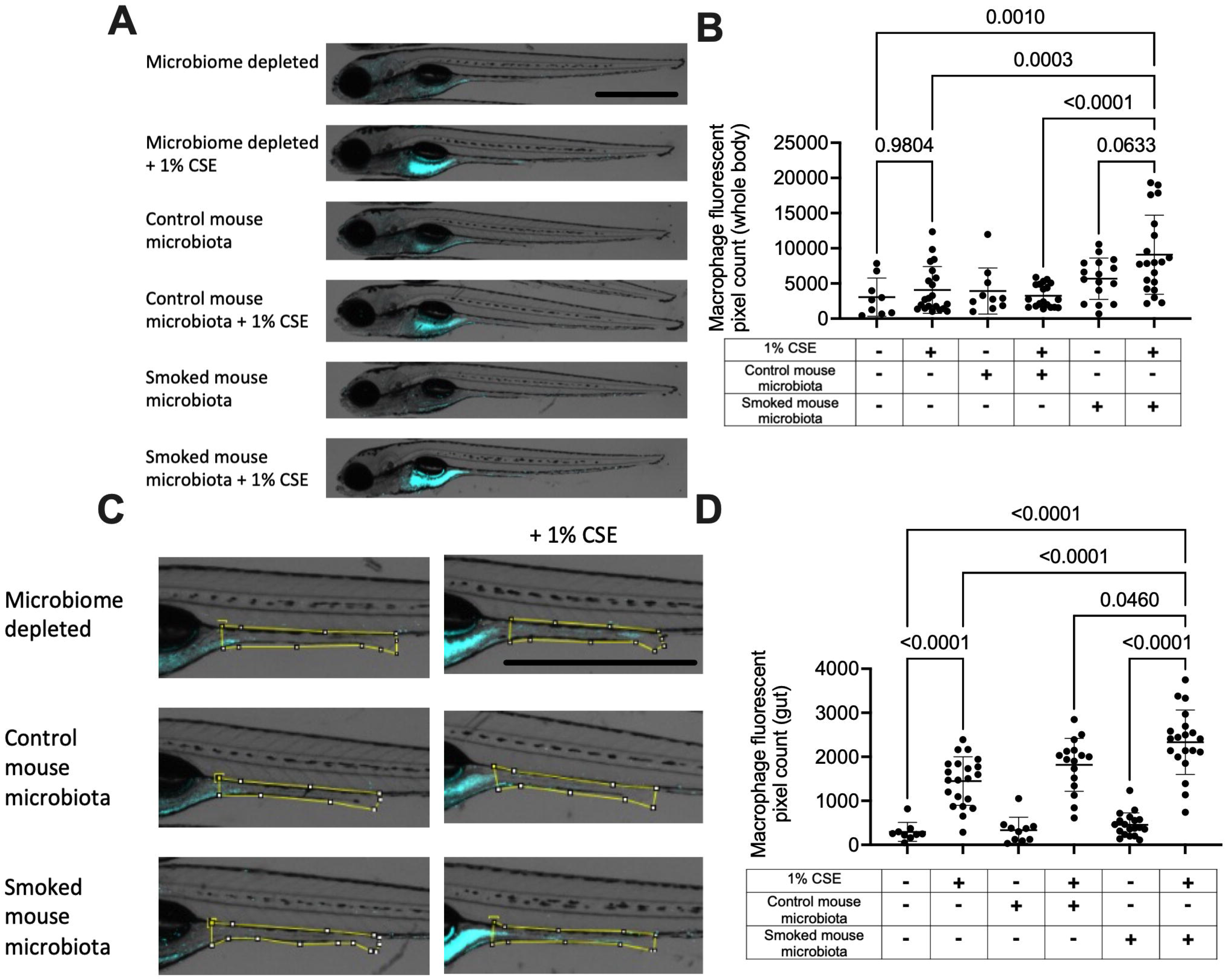
CS-exposed mouse microbiota increases the macrophage inflammatory effects of CSE. A. Representative images of 5 dpf microbiota-depleted *Tg(mfap4:turquoise)*^*xt27*^ larvae exposed to combinations of CS- or control normal air-exposed mouse microbiota with or without 1% CSE from 3 dpf. B. Quantification of total macrophage fluorescent pixel area per larva exposed to combinations of CS- or control normal air-exposed mouse microbiota with or without 1% CSE. Statistical comparison by ANOVA, data are representative of two biological replicates. C. Representative images of the posterior guts of microbiome depleted *Tg(mfap4:turquoise)*^*xt27*^ larvae exposed to combinations of CS- or control normal air-exposed mouse microbiota with or without 1% CSE. Area used for analysis demarcated by yellow box. D. Quantification of gut macrophage fluorescent pixel area per larva exposed to combinations of CS- or control normal air-exposed mouse microbiota with or without 1% CSE. Statistical comparison by ANOVA, data are representative of two biological replicates. Scale bars indicate 1 mm.

### CS-exposed faecal microbiota contains culturable bacteria that do not affect inflammatory responses

To investigate if any individual species of bacteria from CS-exposed mouse microbiota were able to recapitulate the effects of exposure to the bulk microbiota homogenate, we recovered bacteria from the homogenised CS-exposed mouse microbiota-exposed larvae onto rich LB agar. Four species were subcultured to purity by passage on LB agar and genotyped by 16s sequencing as *Acinetobacter radioresistens, Bacteroides vulgatus, Enterobacter cancerogenus, and Stenotrophomonas maltophilia* (Supplementary File). We then screened these individual bacteria by colonising 3 dpf microbiota-depleted larvae in combination with 1% CSE exposure and found no effect on total or gut leukocyte fluorescent area (Figure 6).

**Figure 6.**
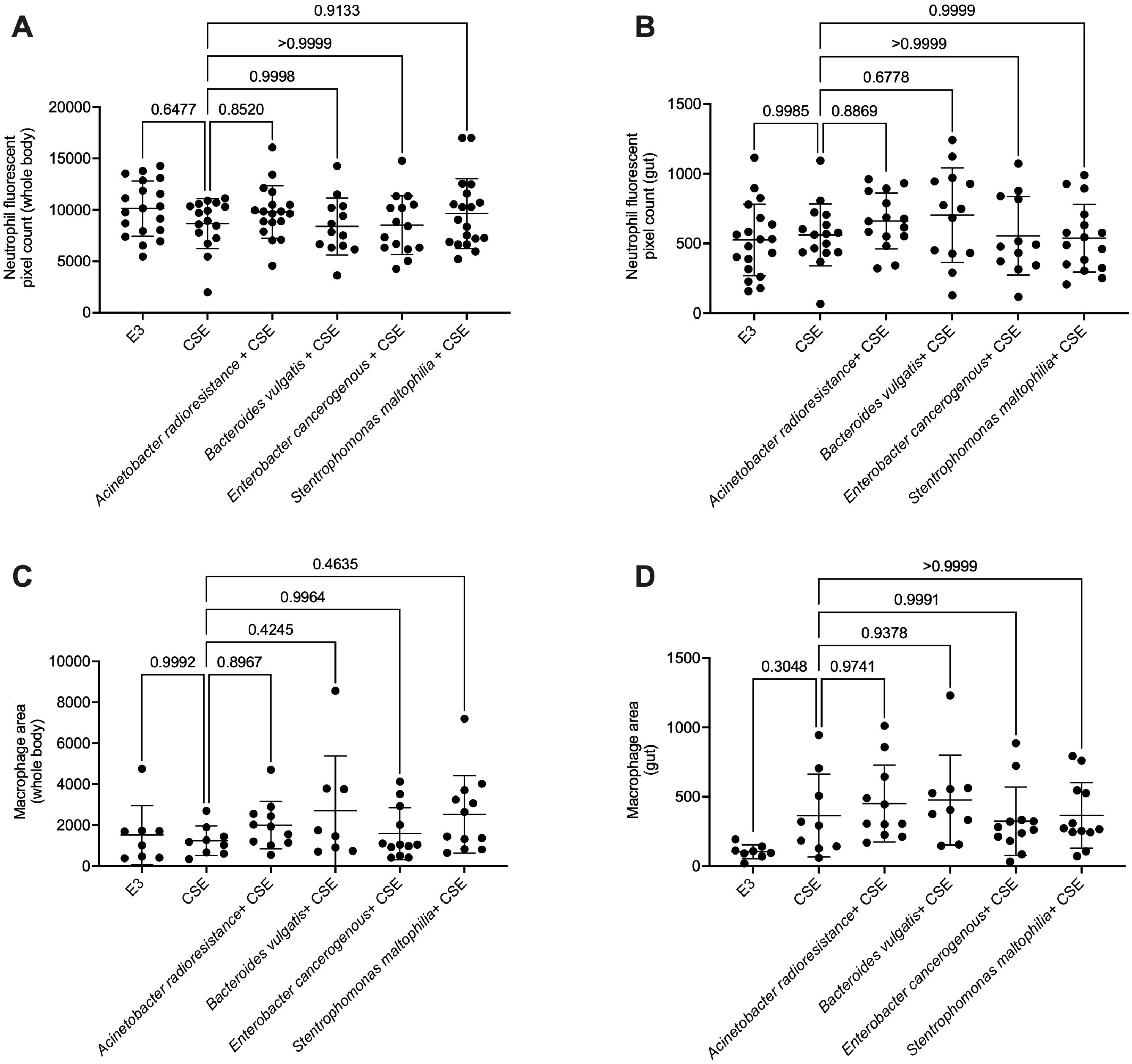
CS-exposed faecal microbiota contain culturable bacteria that do not affect inflammatory responses. A. Quantification of total neutrophil fluorescent pixel area per larva exposed to bacterial isolates with or without 1% CSE. B. Quantification of gut neutrophil fluorescent pixel area per larva exposed to bacterial isolates with or without 1% CSE. C. Quantification of total macrophage fluorescent pixel area per larva exposed to bacterial isolates with or without 1% CSE. D. Quantification of gut macrophage fluorescent pixel area per larva exposed to bacterial isolates with or without 1% CSE. Statistical comparisons by ANOVA.

## Discussion

Here we found the immune system of zebrafish larvae is sensitive to the pro-inflammatory neutrophilic and macrophage effects of CSE. Exposing zebrafish larvae to microbiota derived from the faeces of CS-exposed mice demonstrate the presence of microbes (including culturable *Acinetobacter radioresistens, Bacteroides vulgatus, Enterobacter cancerogenus, and Stenotrophomonas maltophilia*) or metabolites capable of amplifying the inflammatory effects of CSE across species. Together, these data establish zebrafish larvae as a platform for studying the environmental interactions that affect cigarette- and tobacco-induced inflammatory damage.

Our finding that CSE concentrations of >2% are toxic to zebrafish larvae are in line with previous studies by Alvarez et al. who showed larvae can survive in a solution of approximately 1% CSE and Progatzky *et al.* who utilised a 1% CSE challenge with adult zebrafish (19, 21). Exposure of adult zebrafish to CSE by Alvarez *et al.* demonstrated an impairment in post-amputation fin regeneration (19). Our acute wound leukocyte response data adds to this observation by suggesting that increased numbers of wound-associated macrophages may impede physiological regeneration in the presence of CSE.

Our data demonstrate that exposure of zebrafish larvae to 2% CSE causes an expansion of leukocytes (neutrophils and macrophages) and leukocytic inflammation in the gut. Despite the increased leukocyte fluorescent area, we did not observe the mass redistribution of leukocytes to the skin mucosal surfaces as we have previously documented with the enterocolitis-inducing chemical irritants dextran sodium sulfate and 2,4,6-trinitrobenzene sulfonic acid (31, 32). Exposure of zebrafish larvae to a dose of 1% CSE did not cause detectable changes to inflammatory markers in conventionally reared larvae. Our results are comparable to work using adult zebrafish that found only a subtle inflammatory response to 1% CSE exposure in the gill tissue of adult zebrafish with upregulation of inflammatory gene expression but decreased or unchanged immune cell infiltration in the gills depending on the length of exposure (21).

Our system of using a sub-optimal dose of 1% CSE for two days to stimulate a below measurable amount of inflammation proved sensitive to positive modulation by microbiota samples from CS-exposed mouse faeces. Zebrafish have been used to as hosts for trans-species microbiota transplants (33, 34), and our study expands this knowledge by demonstrating sensitivity of recipient zebrafish host inflammation to a microbiota-environment interaction. These proof-of-concept data suggest that zebrafish larvae may be a feasible platform with which human microbiota-CS interactions can be screened to uncover potential missing links in microbiota-environment interactions that appear to predispose some smokers to the development of COPD (4, 9, 11).

The failure of single bacterial species that could be cultured from the guts of zebrafish to recapitulate the pro-inflammatory effects of CS-exposed mouse microbiota in the context of 1% CSE co-exposure suggest either the presence of multiple microbial species and/or other microbiota or factors present in the faecal homogenate. This was not unexpected as the aerobic conditions of the larval zebrafish gut does not facilitate sustained colonisation by some members of the mammalian microbiota (33), and our method of recovering microbes on LB agar in atmospheric conditions will further bias recovery of only species present at a high proportion in the gut of colonised larvae.

Recent mouse studies have demonstrated the importance of the metabolome, including short chain fatty acid output, in susceptibility to smoking-induced pathology (15, 16, 35). Given the lack of butyrate production found in conventionally raised zebrafish larvae (27), we think it is unlikely that exposure to control mouse faecal microbiota produces anti-inflammatory short chain fatty acids as the protective mechanism against 1% CSE exposure compared to the pro-inflammatory CS-exposed mouse faecal microbiota. Instead, we propose that the presence of additional toxins or pro-inflammatory factors either already present or produced by complex mixtures of microbes harvested from the supernatant of faecal homogenates drives inflammatory responses.

The dose response effects of CSE exposure on inflammation in larval zebrafish suggest this platform could be adapted for use as a screening platform to identify positive and negative regulators of inflammation (36). In this study we have demonstrated the utility of a low CSE challenge to identify CS-associated microbiota as factors that increase the susceptibility of zebrafish larvae to CSE-induced inflammation. Follow-up studies would identify other soluble environmental factors that increase the inflammatory potential of CSE. Conversely, a higher dose of 2% CSE challenge could be used as a platform to screen for inhibitors of CSE-induced inflammation. Since we have recently demonstrated the sensitivity of zebrafish larval inflammation to the short chain fatty acid butyrate (27), it would be interesting to determine if high dose CSE-induced inflammation in zebrafish larvae can also be ameliorated by butyrate supplementation.

In summary, we have demonstrated the suitability of zebrafish larvae for studying the inflammatory effects of cigarette smoke-environment interactions and show that the CS-exposed mouse gut microbiota potentiates the inflammatory effects of CSE across host species.

## DATA AVAILABILITY

Raw data is available upon reasonable request to the corresponding authors.

## ACKNOWLEDGEMENTS

We thank other members of the Oehlers lab at the Centenary Institute for their input into this work and maintenance of the zebrafish aquarium; Dr Angela Fontaine and Sydney Cytometry for assistance with imaging; Dr Diana Quan and Ms Trixie Wang for collection of mouse faeces; Dr Kamal Dua for supervision of VM.

**Supplementary File: Identification of isolated bacterial strains**

Top NCBI BLAST hit strain information and sequencing data used for BLAST.

