## Supplementary File for "Exposure to the gut microbiota from cigarette smoke-exposed mice exacerbates cigarette smoke extract-induced inflammation in zebrafish larvae"

**Acinetobacter radioresistens strain WA16094 16S ribosomal RNA gene, partial sequence**

CGCGAGGGGGGCTTACCATGCAAGTCGAGCGGATGAAGGTAGCTTGCTACCGGATTCAGCGGCGGACGGGTGAGTAATGCTTAGGAATCTGCCTATTAGTGGGGGACAACGTTCCGAAAGGAGCGCTAATACCGCATACGTCCTACGGGAGAAAGCAGGGGACCTTTGGGCCTTGCGCTAATAGATGAGCCTAAGTCGGATTAGCTAGTTGGTAGGGTAAAGGCCTACCAAGGCGACGATCTGTAGCGGGTCTGAGAGGATGATCCGCCACACTGGGACTGAGACACGGCCCAGACTCCTACGGGAGGCAGCAGTGGGGAATATTGGACAATGGGGGGAACCCTGATCCAGCCATGCCGCGTGTGTGAAGAAGGCCTTTTGGTTGTAAAGCACTTTAAGCGAGGAGGAGGCTACCTAGATTAATACTTTAGGATAGTGGACGTTACTCGCAGAATAAGCACCGGCTAACTCTGTGCCAGCAGCCGCGGTAATACAGAGGGTGCGAGCGTTAATCGGATTTACTGGGCGTAAAGCGTGCGTAGGCGGCCAATTAAGTCAAATGTGAAATCCCCGAGCTTAACTTGGGAATTGCATTCGATACTGGTTGGCTAGAGTATGGGAGAGGATGGTAGAATTCCAGGTGTAGCGGTGAAATGCGTAGAGATCTGGAGGAATACCGATGGCGAAGGCAGCCATCTGGCCTAATACTGACGCTGAGGTACGAAAGCATGGGGAGCAAACAGGATTAGATACCCTGGTAGTCCATGCCGTAAACGATGTCTACTAGCCGTTGGGGCCCTTGAGGCTTTAGTGGCGCAGCTAACGCGATAAGTAGACCGCCTGGGGAGTACGGTCGCAAGACTAAAACTCAAATGAATTGACGGGGGCCCGCACAAGCGGTGGAGCATGTGGTTTAATTCGATGCAACGCGAAGAACCTTACCTGGCCTTGACATACAGAGAACTTTCCAGAGATGGATTGGTGCCTTCGGGAACTCTGATACAGGTGCTGCATGGCTGTCGTCAGCTCGTGTCGTGAGATGTTGGGTTAAGTCCCGCAACGAGCGCAACCCTTTTCCTTATTTGCCAGCACTTCGGGTGGGAACTTTAAGGATACTGCCAGTGACAAACTGGAGGAAGGCGGGGACGACGTCAAGTCATCATGGCCCTTACGGCAGGGCTACCACGTGCTACATGGTCGGTACAAGGTTGCTACCAGCGATGTGATGCTAATCTCAAAAGCCAATCGNAATCCGGATGGAATCTGCACTCGACCCCTGAATCCGAATCGCTAGAATCCGGAATAAAAAGGCCCGGGGAAAACTTTCCCGGGCCTTGTACCCCCCGCCCCCNNCGGGGGTTTTTTGCCCAAAANAAGGATCTACCCCGNGGAACTTCCCCGGGNNNNAGNAGGGGGNNGGNNNAAAANNCAAANNNNNNNNAAAAAA

**Bacteroides vulgatus strain mpk genome**

GCTTCGCTCCGACGACTACTATGCTAGTCGATCGGTGTCCAGATAGCTTGCTCCCTGCTCCGATGACTTGCTCGACGGTCGGATAATCTATAATGATTGTCTTTTTTAAATCCTCAAAAACAGCCATGAGGACTATGGGGGAAGTCCTGGACAATGTATATATTCAGATATTATAAATGTTCTTTTGGGTTGTTCTGGAAGAGTTAGATCACATGAGATGATATGTCTTGTACCCCATTGCTCATTTTCTATTTCCTTGTATTTTCTTTCCAAACTACCTTCTATAGCCGTGACTCCAAACTTTTCTTGAAGAACCAGTAGTTTGCCTCTTGCACTTTCTGCATCCTTGAAGATGAATTTGTATCCTCCTCCGGTCTTTACTGGTGTTGCACCTTATGCTTTTTCTTTAATTGCTTGTTTTTCAAAGTCTTTTAGTTGTCCCGCAAAAGCTAGAACCTGGATGCTTACAATGTAGTTATCTCCTTTAAGTACTACTATTACGTGAATGTCGCGCGTGTTTCCTGATTGATCTCTTAAAGAACTAATCGCTTCGGTGGGGCTAACGCTTTTATGGATATCTCGAGAGAATGAATTGGATGGTTTCGAGAATTATCGGTTTATCACAACTTATTTCATGCTTCGCGACTCCCTCTTGGGAAATGGAAAGTCTGACTGACTATGAGCACATATCAATTTTCATTTGCTGTCTCTCTAGTCGTTATGACATGGAAGCAAGTACGTGTCTGGAGCTAGAAGACACGAAGAGCCAACTTCGGCGATGTCACAGCCTTCGCTCGGAGCAAGCTTGGGGGGTAAT

**Enterobacter cancerogenus strain YB/JSRM - 03 16S ribosomal RNA gene, partial sequence**

CGCGAGGGCGGCTACACATGCAAGTCGAACGGTAGCACAGAGAGCTTGCTCTCGGGTGACGAGTGGCGGACGGGTGAGTAATGTCTGGGAAACTGCCTGATGGAGGGGGATAACTACTGGAAACGGTAGCTAATACCGCATAACGTCGCAAGACCAAAGAGGGGGACCTTCGGGCCTCTTGCCATCAGATGTGCCCAGATGGGATTAGCTAGTAGGTGGGGTAACGGCTCACCTAGGCGACGATCCCTAGCTGGTCTGAGAGGATGACCAGCCACACTGGAACTGAGACACGGTCCAGACTCCTACGGGAGGCAGCAGTGGGGAATATTGCACAATGGGCGCAAGCCTGATGCAGCCATGCCGCGTGTATGAAGAAGGCCTTCGGGTTGTAAAGTACTTTCAGCGGGGAGGAAGGTGGTGAGGTTAATAACCTCATCGATTGACGTTACCCGCAGAAGAAGCACCGGCTAACTCCGTGCCAGCAGCCGCGGTAATACGGAGGGTGCAAGCGTTAATCGGAATTACTGGGCGTAAAGCGCACGCAGGCGGTCTGTCAAGTCGGATGTGAAATCCCCGGGCTCAACCTGGGAACTGCATTCGAAACTGGCAGGCTAGAGTCTTGTAGAGGGGGGTAGAATTCCAGGTGTAGCGGTGAAATGCGTAGAGATCTGGAGGAATACCGGTGGCGAAGGCGGCCCCCTGGACAAAGACTGACGCTCAGGTGCGAAAGCGTGGGGAGCAAACAGGATTAGATACCCTGGTAGTCCACGCCGTAAACGATGTCGACTTGGAGGTTGTGCCCTTGAGGCGTGGCTTCCGGAGCTAACGCGTTAAGTCGACCGCCTGGGGAGTACGGCCGCAAGGTTAAAACTCAAATGAATTGACGGGGGCCCGCACAAGCGGTGGAGCATGTGGTTTAATTCGATGCAACGCGAAGAACCTTACCTACTCTTGACATCCAGAGAACTTACCAGAGATGCATTGGTGCCTTCGGGAACTCTGAGACAGGTGCTGCATGGCTGTCGTCAGCTCGTGTTGTGAAATGTTGGGTTAAGTCCCGCAACGAGCGCAACCCTTATCCTTTGTTGCCAGCGGTTAGGCCGGGAACTCAAAGGAGACTGCCAGTGATAACTGGAGGAAGGTGGGGATGACGTCAAGTCATCATGGCCCTTACGAGTAGGGCTACCACGTGCTACATGGCGCATACAAGAAGAACGACCTCGNGAGAGNANCGGACTCCNAAAGGGGGCNNAAATCCGGATGGAATCGGCACTCCACCCCTGAAGTCGAATCGCTGAAATCGGGATCAAAGGCCCGGGGAAAACTTCCCGGCCTGGACCCCCCGGCCCCCCCGGGGGGGGGGGGGAAAAAAAAGNNTTTCCTCGGGGCTNTTNTTAGGGNNNGGGGGGGGGNNGGGGGGGGGGGAGNAAAAAAAAAAG

**Stenotrophomonas maltophilia strain yy01 16S ribosomal RNA gene, partial sequence**

CGCGGGGCGTAGGCTANCATGCAAGTCGAACGGCAGCACAGGAGAGCTTGCTCTCTGGGTGGCGAGTGGCGGACGGGTGAGGAATACATCGGAATCTACTTTTTCGTGGGGGATAACGTAGGGAAACTTACGCTAATACCGCATACGACCTACGGGTGAAAGCAGGGGATCTTCGGACCTTGCGCGATTGAATGAGCCGATGTCGGATTAGCTAGTTGGCGGGGTAAAGGCCCACCAAGGCGACGATCCGTAGCTGGTCTGAGAGGATGATCAGCCACACTGGAACTGAGACACGGTCCAGACTCCTACGGGAGGCAGCAGTGGGGAATATTGGACAATGGGCGCAAGCCTGATCCAGCCATACCGCGTGGGTGAAGAAGGCCTTCGGGTTGTAAAGCCCTTTTGTTGGGAAAGAAATCCAGCTGGTTAATACCCGGTTGGGATGACGGTACCCAAAGAATAAGCACCGGCTAACTTCGTGCCAGCAGCCGCGGTAATACGAAGGGTGCAAGCGTTACTCGGAATTACTGGGCGTAAAGCGTGCGTAGGTGGTCGTTTAAGTCCGTTGTGAAAGCCCTGGGCTCAACCTGGGAACTGCAGTGGATACTGGACGACTAGAGTGTGGTAGAGGGTAGCGGAATTCCTGGTGTAGCAGTGAAATGCGTAGAGATCAGGAGGAACATCCATGGCGAAGGCAGCTACCTGGACCAACACTGACACTGAGGCACGAAAGCGTGGGGAGCAAACAGGATTAGATACCCTGGTAGTCCACGCCCTAAACGATGCGAACTGGATGTTGGGTGCAATTTGGCACGCAGTATCGAAGCTAACGCGTTAAGTTCGCCGCCTGGGGAGTACGGTCGCAAGACTGAAACTCAAAGGAATTGACGGGGGCCCGCACAAGCGGTGGAGTATGTGGTTTAATTCGATGCAACGCGAAGAACCTTACCTGGCCTTGACATGTCGAGAACTTTCCAGAGATGGATTGGTGCCTTCGGGAACTCGAACACAGGTGCTGCATGGCTGTCGTCAGCTCGTGTCGTGAGATGTTGGGTTAAGTCCCGCAACGAGCGCAACCCTTGTCCTTAGTTGCCAGCACGTAATGGTGGGAACTCTAAGGAGACCGCCGGTGACAAACCGGAGGAAGGTGGGGATGACGTCAAGTCATCATGGCCCTTACGGCCAGGGCTACCCCGTACTACAATGGTAGGGANGAAGGGCTGCAAGCCGGCGACGGTAAGCCATCCNAAAACCCTATTCCAGTCCGGATGGNGTCGGCACTCCACTCCTGAAATCGGAATCCCTAGAAATCCCAAAAAACCATTGGTGCCGGGAAAAACNTCCCGGGCCNNNNACCCCCCCCCCCCCCCCCCGGGGAATTTTTTGCNCAAAAAAGGGNNNTAACCCTNNNNNGGNNNGGCCGGGGGCGAAGGANNNNNGGGAGGAAGAANAAAAANCANNAAAAAAAAAAANAAAAA
